# Optimization of 4D Combined Angiography and Perfusion using Radial Imaging and Arterial Spin Labeling

**DOI:** 10.1101/2022.07.13.499856

**Authors:** Thomas W. Okell, Mark Chiew

## Abstract

**Purpose:** To extend and optimize a non-contrast MRI technique to obtain whole head 4D (time-resolved 3D) angiographic and perfusion images from a single scan.

**Methods:** 4D combined angiography and perfusion using radial imaging and arterial spin labeling (CAPRIA) uses pseudocontinuous labeling with a 3D golden ratio (“koosh ball”) readout to continuously image the blood water as it travels through the arterial system and exchanges into the tissue. High spatial/temporal resolution angiograms and low spatial/temporal resolution perfusion images can be flexibly reconstructed from the same raw k-space data at any timepoint within the readout. A constant flip angle (CFA) and a quadratic variable flip angle (VFA) excitation schedule were optimized through simulations and tested in healthy volunteers. A conventional sensitivity encoding (SENSE) reconstruction was compared against a locally low rank (LLR) reconstruction, which leverages spatiotemporal correlations to improve reconstruction quality. Differences in image quality were assessed through split-scan repeatability.

**Results:** The optimized VFA schedule (2-9°) reduced initial signal attenuation whilst boosting the signal at later timepoints, resulting in a significant (p < 0.001) improvement in image quality (up to 84%), particularly for the lower SNR perfusion images. The LLR reconstruction provided effective denoising without biasing the signal timecourses, significantly improving angiographic and perfusion imaging repeatability (up to 143%, p < 0.001) and enabling the clear depiction of small distal vessels and late tissue perfusion.

**Conclusion:** 4D CAPRIA can be optimized using a VFA schedule and LLR reconstruction to yield whole head 4D angiograms and perfusion images from a single scan.

## Introduction

The ability to visualize blood flow to the brain, both through the arterial system (angiography) and at the level of the tissue (perfusion), is crucial in many cerebrovascular diseases, including stroke and arteriovenous malformation^1^. X-ray based methods require the administration of an exogenous contrast agent and exposure to ionizing radiation. Contrast-enhanced magnetic resonance imaging (MRI) methods avoid ionizing radiation, but often have limited spatiotemporal resolution due to the necessity to image the first passage of the bolus^2^, and there are concerns about the use of Gadolinium-based contrast agents in patients with kidney disease^3^, as well as accumulation in the brain^4^. A non-contrast alternative is therefore desirable.

Arterial spin labeling (ASL) is an MRI-based non-contrast method capable of generating both angiograms^5–7^ and maps of tissue perfusion^8,9^. However, the high spatial and temporal resolution required for angiography makes time-resolved 3D ASL angiograms slow to acquire in a conventional manner^2^, whilst ASL perfusion images require the acquisition of many averages to improve the signal-to-noise ratio (SNR)^10^. Therefore, obtaining both angiographic and perfusion information using ASL within a busy clinical protocol is often challenging.

Recently, a few methods have been proposed to address this problem by acquiring angiograms and perfusion maps simultaneously from a single ASL acquisition, thereby increasing the time-efficiency. This has involved either the use of two separate readout modules for angiography and perfusion imaging^11^, or the use of a single golden ratio^12^ radial readout^13,14^. This latter approach allows flexibility in the spatial and temporal resolution of the reconstructed images, thereby allowing high spatial/temporal resolution angiograms and lower spatial/temporal resolution perfusion maps to be generated at multiple time points after the labeling period from the same raw k-space data. However, this approach has thus far only been demonstrated using single or multi-slice 2D readouts, limiting the spatial coverage and SNR achievable.

In this work, we extend and optimize the image quality of one of these methods, Combined Angiography and Perfusion using Radial Imaging and ASL (CAPRIA)^13^, by: 1) utilizing multi-dimensional golden means to allow fully isotropic 4D (time-resolved 3D) imaging of the whole head; 2) modulating the ASL signal using a variable flip angle readout, mitigating signal attenuation due to the imaging pulses; and 3) applying an advanced image reconstruction technique to leverage spatiotemporal signal correlations and minimize noise propagation in these undersampled acquisitions. Here we focus on optimizing qualitative angiography and perfusion imaging. The extraction of quantitative physiological parameters from this kind of data will be explored in future work. This study builds upon work previously presented in abstract form^15–17^.

## Methods

### Pulse Sequence Design

A schematic of the 4D CAPRIA sequence is shown in Figure 1. The preparation period is identical to the original approach^13^, consisting of a water suppression enhanced through T_1_ effects (WET) pre-saturation module^18^ for background suppression and a 1400 ms duration balanced pseudocontinuous (PCASL)^19^ pulse train to label blood flowing through a plane in the neck (positioned as per previous studies^20,21^).

**Figure 1:**
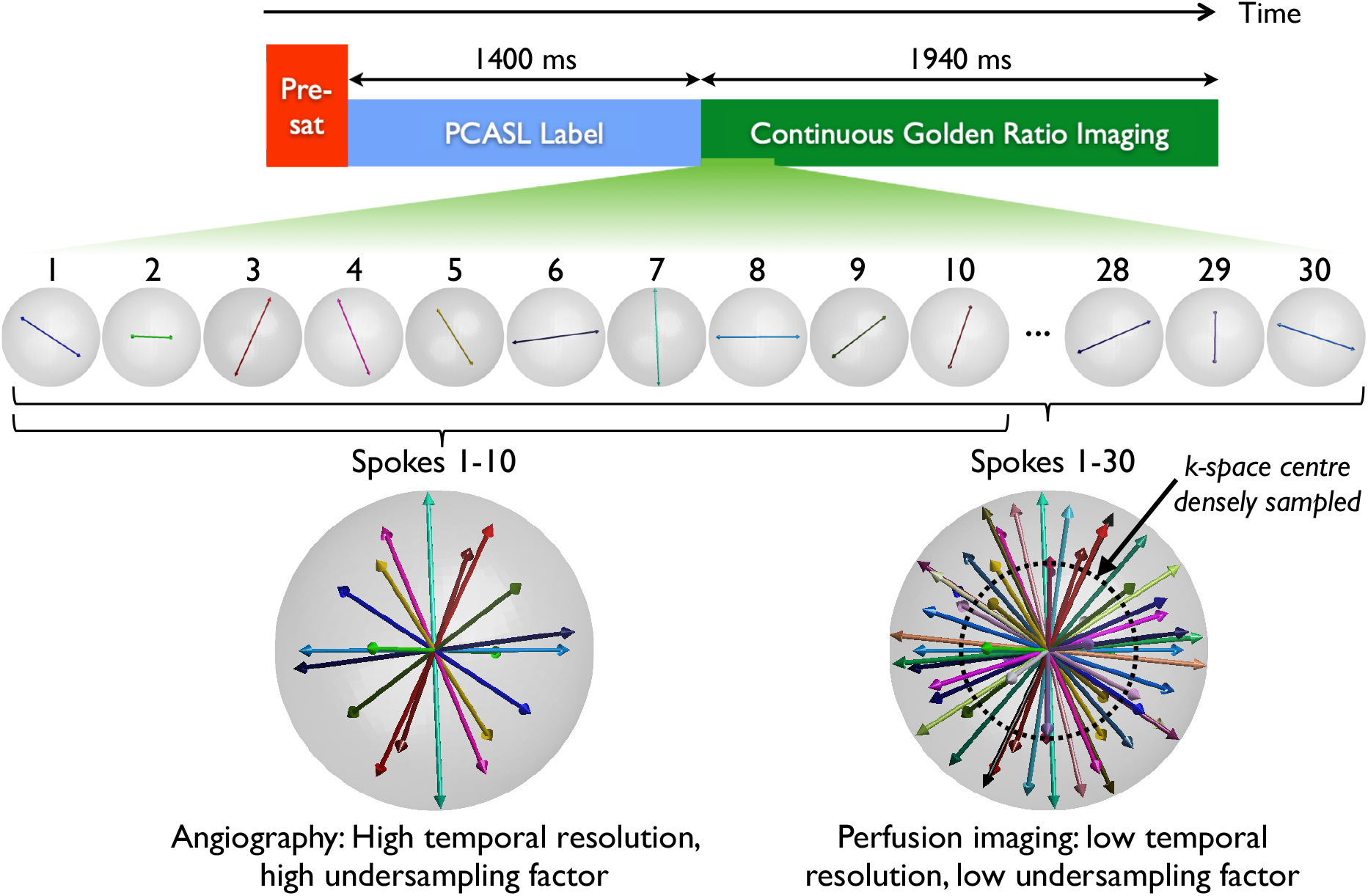
4D CAPRIA sequence schematic. A WET pre-saturation module (pre-sat) is followed by PCASL labeling and then continuous 3D golden ratio imaging, providing flexibility for the image reconstruction. In this simple example, sets of 10 spokes can be used to reconstruct angiographic images with a high temporal resolution (small temporal window) at any time point within the readout period. This results in a high undersampling factor, but this can be tolerated for the sparse, high SNR angiographic signal. The same raw k-space data can be reconstructed at a lower temporal resolution (larger temporal window, 30 spokes used here), which is better suited to the slower dynamics of perfusion imaging. This results in a lower undersampling factor, necessary for the low SNR and less sparse perfusion signal. Using just the densely sampled centre of k-space, lower spatial resolution perfusion images can be reconstructed, further reducing the undersampling factor.

This is followed by a continuous 3D golden ratio radial (“koosh ball”) spoiled gradient echo readout, where a single line (“spoke”) of data going through the k-space origin is acquired following each of a series of non-selective excitation pulses. The polar and azimuthal angles of each spoke are calculated according to the multi-dimensional golden means^22^, which is a 3D generalization of the original golden ratio approach^12^. The azimuthal *(ϕ)* and polar *(θ)* angles of the *m*^th^ radial spoke are:

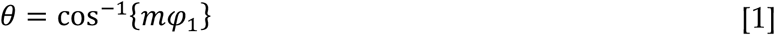

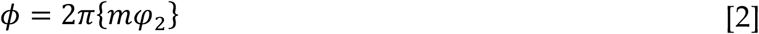

Where *φ*_*1*_ = 0.4656…, *φ*_*2*_ = 0.6823… and the curly braces {..} denote taking the fractional part (i.e. modulus 1). This approach retains the same appealing property that any set of contiguously acquired spokes gives an approximately uniform coverage of k-space, meaning that the temporal window used for image reconstruction (and therefore the temporal resolution, undersampling factor and post-labeling delay [PLD]) can be arbitrarily chosen retrospectively.

As illustrated in Figure 1, the golden ratio trajectory allows angiographic images to be reconstructed with a small temporal window (high temporal resolution), which is required to capture the rapid flow of blood through the arterial system. This results in a high undersampling factor, but the spatially sparse and high SNR nature of the angiographic signal mean this can be tolerated^13,23,24^. Due to T_1_ decay and the dilution of the labeled blood, the ASL signal is considerably weaker by the time it reaches the tissue. However, the dynamics of this perfusion signal are also slower, so a broader temporal window can be used to reconstruct perfusion images from the same raw k-space data. In addition, the centre of k-space is much more densely sampled than the periphery, so lower spatial resolution images, as are typically used for perfusion imaging, can be reconstructed with a much lower undersampling factor. The ability to reconstruct angiographic or perfusion-like images at any timepoint within the readout period retrospectively gives a great degree of flexibility to adapt to the hemodynamics of any individual subject.

In practice, data combined across many ASL preparation periods are required to achieve sufficient sampling. As previously^13^, we define a maximum temporal window, *t*_*max*_, at which data will be reconstructed, which corresponds to M radial spokes (*M = t*_*max*_ */ TR*). The golden ratio spoke counter, *m*, for the *i*^*th*^ spoke acquired after the *n*^*th*^ ASL preparation is then:

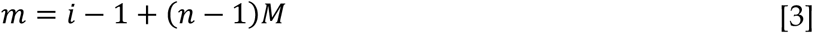

In order to minimize subtraction artifacts due to factors such as scanner drift or motion the acquisition was interleaved such that the time between the same k-space spokes being acquired for the ASL label and control conditions was as short as possible.

A readout partial Fourier (asymmetric echo) factor of 0.79 was used in order to reduce the echo time and minimize flow-induced dephasing effects. However, Eq. [1] results in each spoke starting on the surface of one hemisphere (0 < *θ* < π/2), so readout partial Fourier would lead asymmetric sampling of k-space. Therefore, the direction of every other spoke was reversed to give a more even distribution, i.e. for odd values of *m*:

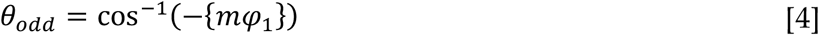

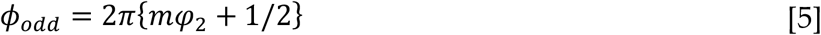

Finally, two possible flip angles schedules for the excitation pulses were investigated: a) a constant flip angle (CFA) approach and b) a variable flip angle (VFA) approach, in which the flip angle of the *i*^*th*^ pulse in the readout, α_i_, increases quadratically from α_1_ at the first pulse to α_N_ at the final (*N*^*th*^) pulse as follows^25^:

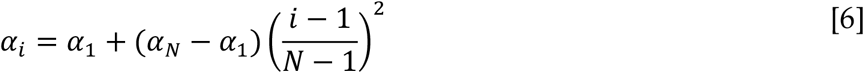

This could help to minimize signal attenuation at early timepoints whilst boosting the signal at later timepoints, which is anticipated to be particularly beneficial for the CAPRIA perfusion signal that relies on the accumulation of labeled blood water over time.

### Parameter optimization simulations

It was necessary to reoptimize the readout parameters (the repetition time of the excitation pulses, TR, and their flip angles, α_i_) for 4D CAPRIA in order to balance the need for fast acquisitions, low undersampling factors and strong initial signal against signal attenuation effects. Based on the parameters required for *in vivo* scanning (Table 1), a TR of 9 ms was chosen to minimize signal attenuation whilst keeping the undersampling factors within reasonable limits.

**Table 1:**
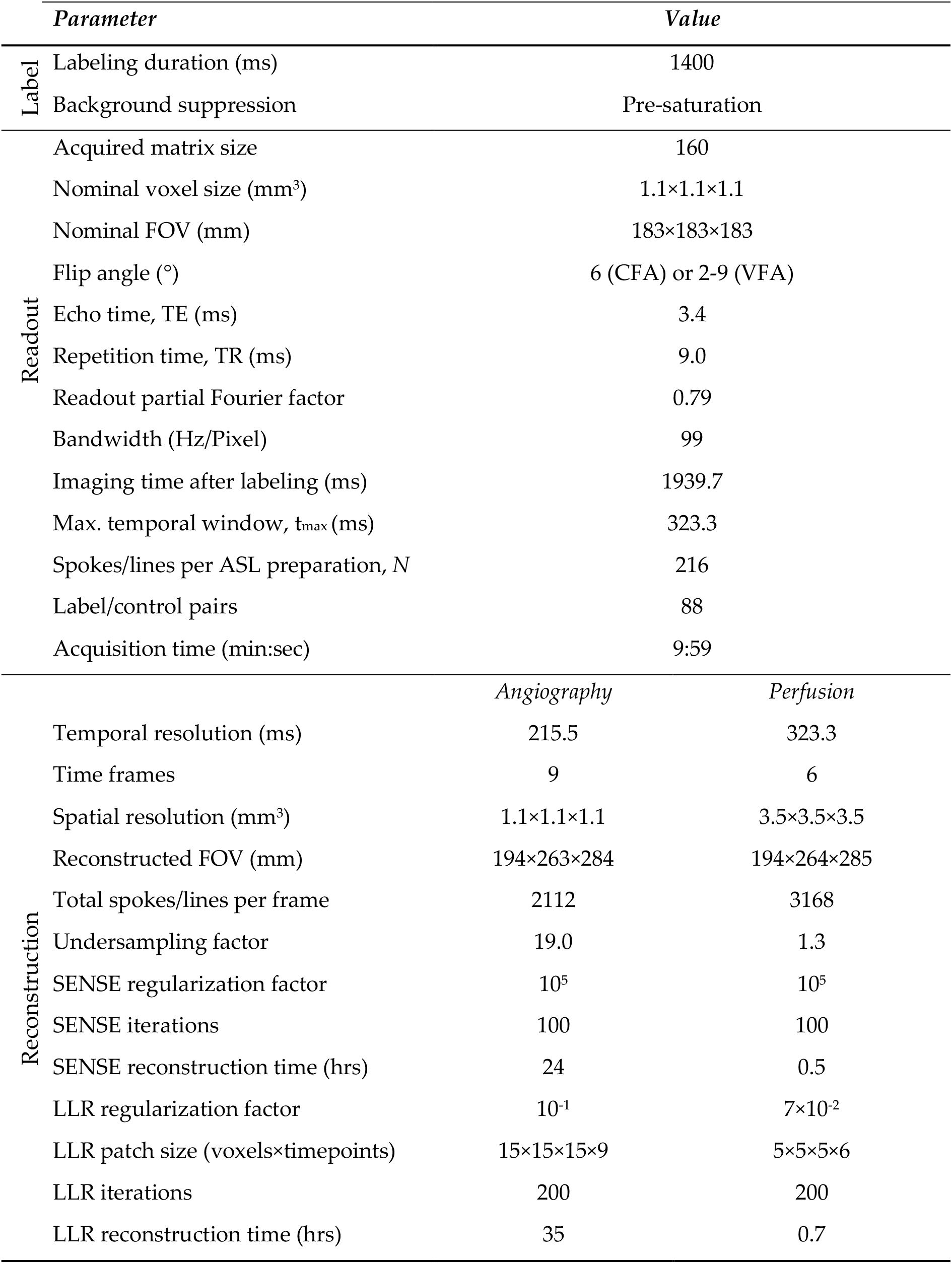
Imaging and reconstruction parameters

CAPRIA acquisition parameters were optimized by simulating the relative strength of the angiographic and perfusion signals across a range of possible parameter choices^13^. The angiographic signal was evaluated using a kinetic model for PCASL angiography^26^, assuming the labeled blood experiences all of the RF excitation pulses due to their non-selective nature. However, the RF attenuation factor, *R*_*i*_, applied to the signal acquired after the *i*^*th*^ RF pulse must be generalized to account for a VFA schedule:

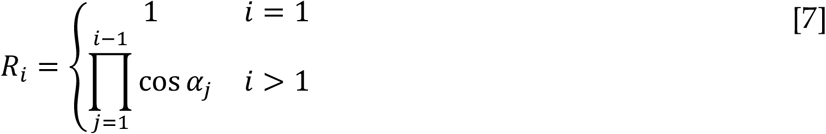

In addition, the VFA effect on the magnitude of the excited transverse magnetization must be also be accounted for. For this study, we also ignore the effects of dispersion to give a final simplified angiographic model:

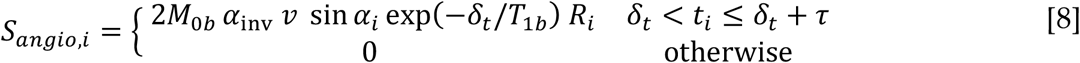

Where *t*_*i*_ is the time of excitation pulse *i* relative to the start of PCASL labeling, *δ*_*t*_ is the macrovascular transit time, *τ* is the PCASL labeling duration, *v* is the fractional macrovascular blood volume and we have expressed the scaling factor using the equilibrium magnetization of blood, *M*_*0b*_, and PCASL inversion efficiency, α_inv_.

Similarly, the CAPRIA perfusion signal^13^, *S*_*perf,i*_, after the *i*^*th*^ excitation pulse, must also be modified:

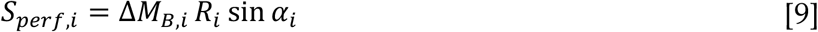

where Δ*M*_*B,i*_ is the Buxton model for the (P)CASL perfusion signal^27^.

The angiographic and perfusion signals were simulated for a range of flip angle schedules (α1 and αN) across a range of physiological parameters. Since blood volume and blood flow (to a first approximation) only scale the angiographic and perfusion signals, respectively, it is not necessary to optimize over these parameters explicitly^28^, leaving only the macrovascular transit time, *δ*_*t*_, and the tissue transit time, *Δt*, to optimize over. Here, we optimized for 0.2 s < *δ*_*t*_ < 1 s and 0.5 s < *Δt* < 2 s.

To compare different acquisition parameters, the mean angiographic signal was calculated during the time that labeled blood was present (i.e. *δ*_*t*_ ≤ *t*_*i*_ < *τ*+ *δ*_*t*_). Similarly, the mean perfusion signal was calculated after all blood water had reached the voxel (*t* ≥ *τ* + *Δt*) when the perfusion signal is approximately proportional to cerebral blood flow^27^. These mean signals were then averaged across all physiological parameters and normalized to the maximum value to give a scalar performance metric for angiography, 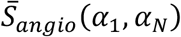 and perfusion,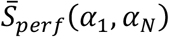. A combined measure,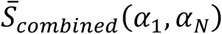, was also generated by a weighted sum of these two and renormalized:

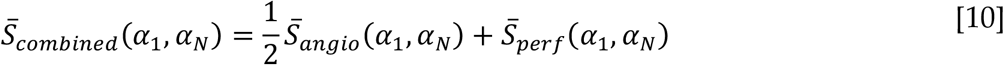

The arbitrary weighting factor of ½ was chosen on the assumption that the perfusion signal is considerably lower SNR and therefore should carry a higher weight.

### Subjects and scan protocol

To demonstrate 4D CAPRIA and compare CFA and VFA approaches, four healthy volunteers (two female, age range 26 – 39) were scanned under a technical development protocol agreed by local ethics and institutional committees on a 3T Verio scanner (Siemens Healthineers, Erlangen, Germany) using a 32-channel head coil. A short (29s) time-of-flight angiogram was acquired to help position the ASL labeling plane^21^, along with a short (2.5 min) T1-weighted structural image for anatomical reference (MP-RAGE^29^, 1.8 mm voxel size, 900 ms TI).

1.1 mm isotropic CFA and VFA 4D CAPRIA scans covering the head and neck were then performed using optimal flip angles derived from simulations (CFA: α = 6°; VFA: α1 = 2°, αN = 9°). Other acquisition and reconstruction parameters are listed in Table 1.

### Image reconstruction

From each raw CAPRIA k-space dataset, two sets of images were reconstructed: 1) angiographic images with high spatial/temporal resolution (1.1mm isotropic, 216 ms, undersampling factor = 19.0); and 2) perfusion images with reduced spatial/temporal resolution (3.5 mm isotropic, 323 ms, undersampling factor = 1.3) to improve the SNR. To minimize computational requirements, perfusion images were reconstructed directly at a lower matrix size using data from the central region of k-space.

Coil sensitivities were estimated by combining data across all time frames, averaging the label/control data and applying a Hann filter to minimize signal aliasing arising from undersampling. The non-uniform fast Fourier transform (NUFFT)^30,31^ operator was then used with a short conjugate gradient iterative reconstruction to produce one image per coil element at the same resolution intended for reconstruction prior to coil sensitivity estimation^32^ and masking. Coil compression from 32 to 12 channels was performed to reduce computational burden^33^.

The phase of the same k-space spokes acquired in label and control conditions were aligned^34^ prior to complex subtraction in k-space to minimize any phase inconsistencies. The ASL difference images were then reconstructed directly using two different approaches: 1) conventional iterative conjugate gradient sensitivity encoding (SENSE)^35^with L2 regularization; and 2) a locally low rank (LLR) approach^36^, using cycle spinning^37^ and the proximal optimized gradient method^38^, to leverage spatiotemporal correlations in the patterns of blood flow.

All reconstruction stages were run using MATLAB 2017b (Mathworks, Natick, MA), incorporating the IRT NUFFT^31^, on a CentOS cluster. Regularization factors and LLR patch sizes were chosen empirically based on preliminary experiments (see Table 1). The absolute value of the complex angiographic and perfusion signals were taken prior to visualization and further analysis.

### Repeatability

Conventional SNR measurements are challenging in this setting due to signal aliasing, spatially varying noise and the use of a non-linear reconstruction method (LLR). Therefore, as an alternative measure of image quality, a repeatability metric was calculated to give an indication of signal stability. The raw k-space data was split into two halves, each half used to reconstruct angiograms and perfusion images, and the Pearson correlation coefficient between the two halves calculated at each timepoint within a spatial mask. This mask excluded large regions of noise-only background which could bias the correlation metric.

For perfusion imaging, a whole brain mask was used, generated from the T1-weighted structural image (using fsl_anat^39^) and linearly registered^40^ to the CAPRIA data. For angiography, a dilated vessel mask was created to include all the major vessels and some background by: averaging CFA and VFA angiograms reconstructed with SENSE, taking the temporal mean, multiplying by the brain mask, thresholding at 0.7 × 99.9^th^ percentile image intensity, then dilating this mask using a spherical kernel (radius 3.5 mm). An example mask is shown in Supporting Information Figure S1.

Prior to performing statistical analyses, these correlation coefficients, *r*, were Fisher transformed^41^ as follows:

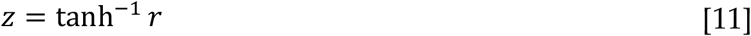

A multi-way ANOVA was performed to assess the statistical significance of differences in *z* between CFA and VFA schedules or SENSE and LLR reconstructions, with subject number and PLD as additional factors.

## Results

### Parameter optimization simulations

Example CFA and VFA flip angle schedules and the resulting simulated angiographic/perfusion signals are shown in Figure 2A-C. As expected, the use of a CFA schedule results in the attenuation of both the angiographic and perfusion signals over time, but it is particularly problematic in the perfusion case due to the later blood arrival times. In contrast, the VFA schedule results in a smaller, but more consistent, angiographic signal that increases slightly over time before the labeled blood water washes out. However, it has a double benefit for the perfusion signal: reduced attenuation at early timepoints means a larger perfusion signal can accumulate within the voxel, but also the amount of signal generated by the excitation pulses at later timepoints is higher. In the case shown (*Δt* = 1.5 s), this resulted in a 2.3x larger perfusion signal at the end of the readout.

**Figure 2:**
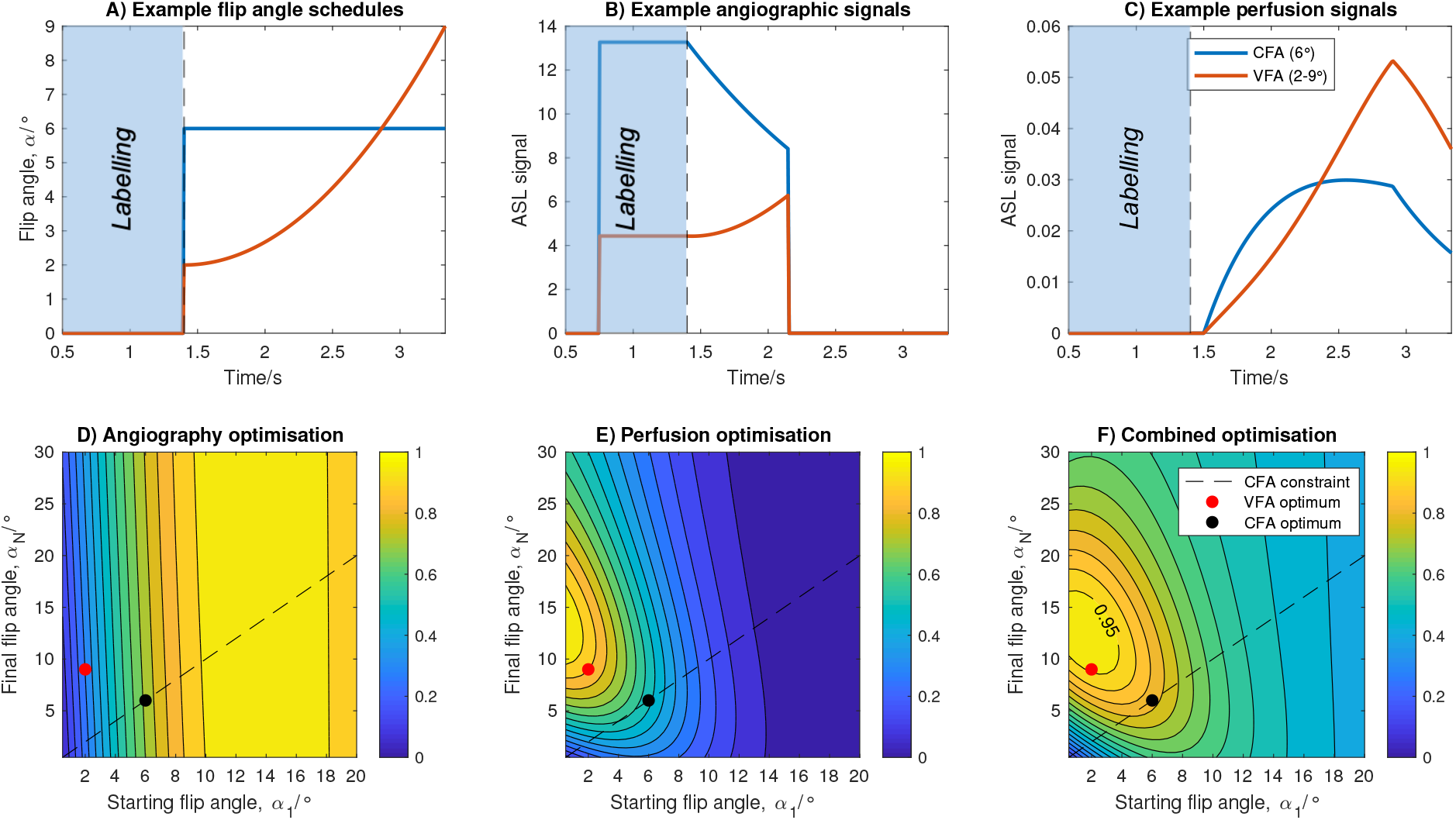
Flip angle schedules (A), simulated angiographic signals (B) and simulated perfusion signals (C) for one example constant flip angle schedule (CFA = 6°) and one quadratically varying flip angle schedule (VFA = 2-9°), demonstrating the potential for considerable signal gains in the perfusion images using VFA (these plots assume transit times of 0.75 s and 1.5 s to the arterial and perfused tissue voxels, respectively). Optimization of the flip angle parameters to maximize signal strength over a range of physiological parameters is shown for angiography (D), perfusion (E) and a combination (weighted sum) of the two (F). Each contour plot is normalized, the constraint corresponding to a constant flip angle shown as a dashed line, and the final chosen optimum CFA and VFA parameters plotted as black and red circles, respectively.

The results of optimizing the flip angle schedules across a range of physiological parameters are shown in Figure 2D-F. The angiographic optimization favors larger starting flip angles with less dependence on the final flip angle, since most of the signal is present at early timepoints. Conversely, the perfusion optimization favors low starting flip angles and higher final flip angles. The combined optimization metric had a broad peak, so the (*α*_*1*_, *α*_*N*_) combination that was within 95% of the peak value but closest to the origin was chosen for the remainder of the study (2-9°) to minimize the signal attenuation effect on late arriving blood. The CFA schedule is constrained to lie on the line *α*_*1*_ = *α*_*N*_, which gives considerably poorer predicted performance for the perfusion and combined optimization metrics than the VFA approach. The CFA flip angle which optimized the combined metric was 6° and this was used for the remainder of this study.

### In vivo results

Example 4D CAPRIA data acquired with a CFA schedule and reconstructed with SENSE are shown in Figure 3. Despite the high undersampling factor (R = 19) the reconstructed angiograms clearly show the flow of blood through the arterial system. Note that, due to the long labeling duration (1.4 s), the first frame shows much of the vasculature filled with labeled blood, but inflow visualization can also be achieved retrospectively (see discussion). There was some minor signal loss in proximal vessels, most likely due to flow-induced dephasing where the blood is changing direction rapidly. Reduced signal intensity at later timepoints is also evident due to the attenuating effect of the CFA excitation pulses.

**Figure 3:**
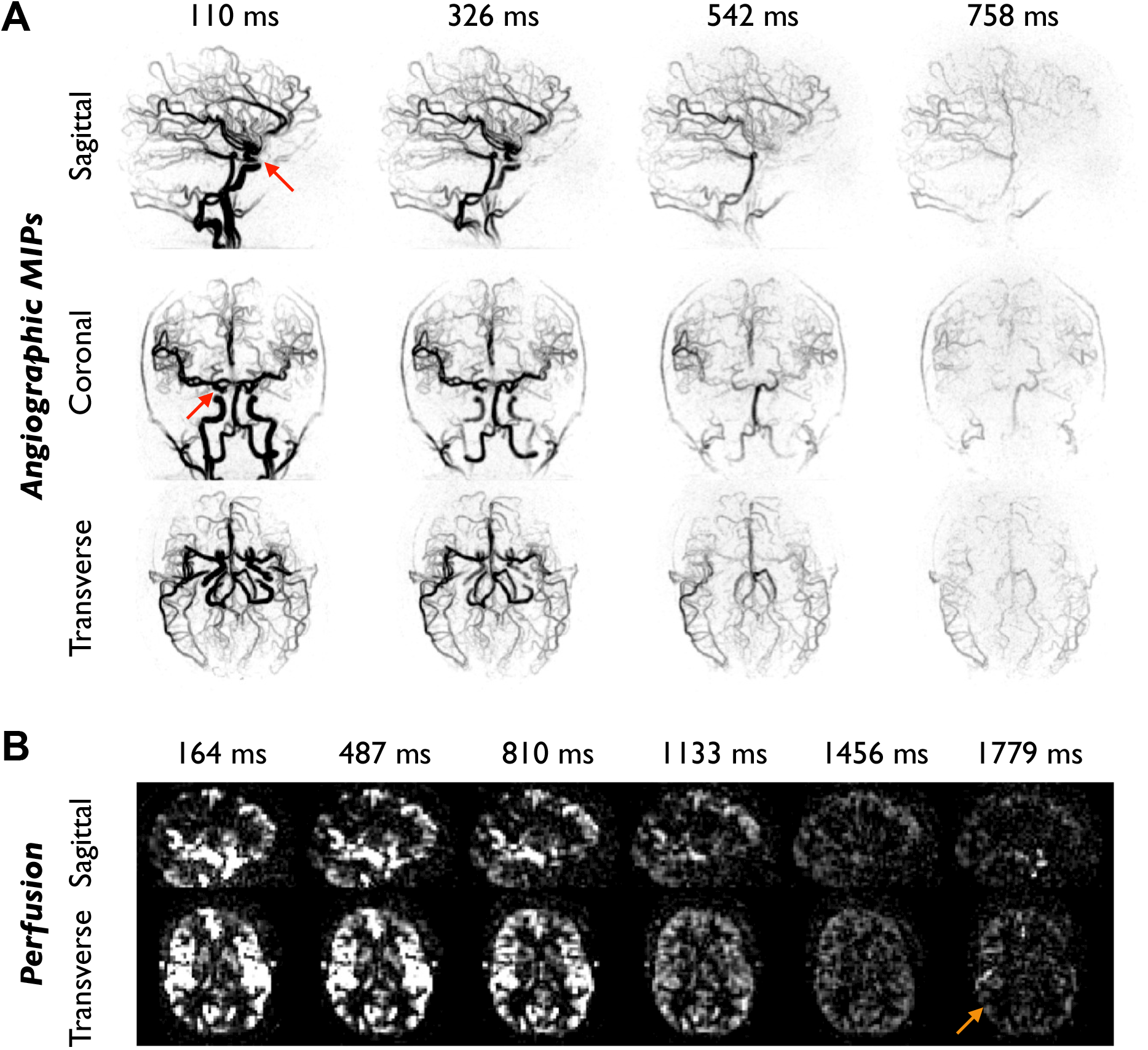
Example CAPRIA SENSE reconstructions in subject 1 using a constant flip angle (CFA) schedule: A) Angiography maximum intensity projections in inverted greyscale at selected PLDs demonstrate good vessel visualization and show the dynamic passage of the bolus through the vasculature. Some minor loss of signal is noted where the blood is changing direction rapidly (red arrows). B) Example sagittal and transverse slices of the perfusion images reconstructed from the same raw dataset, showing mainly macrovascular signal at early PLDs but perfusion of the tissue at later PLDs. However, SNR at later timepoints is low, limiting the ability to visualize tissue perfusion clearly (orange arrow). PLDs are indicated above each subfigure.

Within the perfusion images reconstructed from the same raw k-space data, blood was well visualized at early timepoints, showing ASL signal within the large vessels and in tissue regions where blood arrives quickly (e.g. some deep grey matter structures). However, the majority of the tissue is not perfused until later timepoints when the attenuation effect has become more significant. This effect, combined with the more dispersed blood signal and greater T1 decay, causes the perfusion image SNR to be very low.

Figure 4 shows angiographic and perfusion SENSE reconstructions from the same subject as Figure 3, but this time acquired with a VFA schedule. Despite the low initial flip angle (2°), the angiographic images at early timepoints are qualitatively similar to those produced by the CFA approach. The signal does not reduce over time and visualization of distal blood vessels at later timepoints is improved. As predicted from simulations, the increase in signal at later timepoints achieved with the VFA approach means tissue perfusion is much more clearly visualized. In addition, the spoiled gradient echo 3D radial readout results in isotropic spatial resolution with no through-slice blurring, distortion or dropout artifacts, which are common in other ASL perfusion imaging approaches^10^.

**Figure 4:**
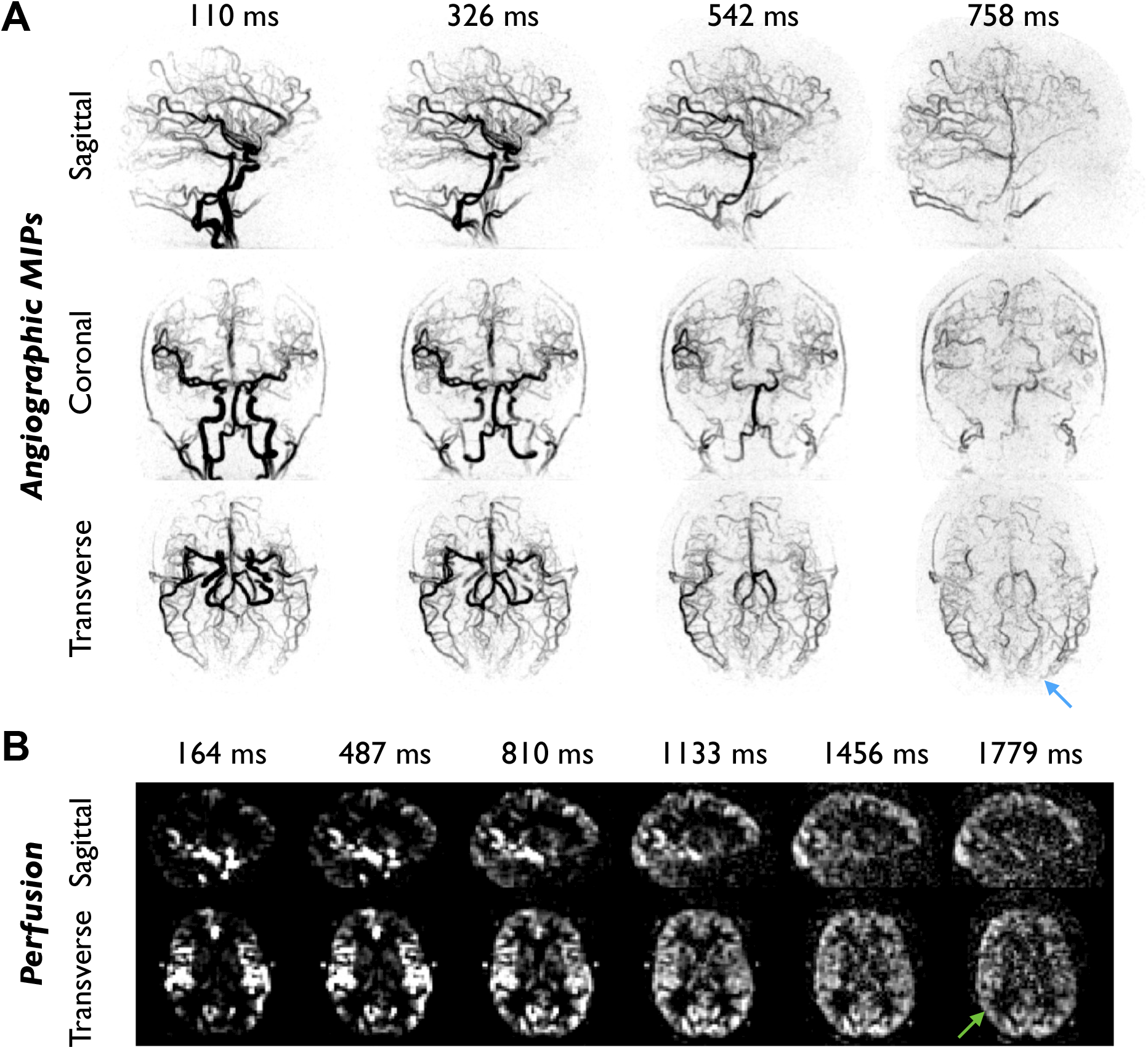
Example CAPRIA SENSE data from the same subject shown in Figure 3 (subject 1), but this time acquired with a variable flip angle (VFA) scheme: A) Despite the considerably lower initial flip angle, angiographic maximum intensity projections at early PLDs show comparable image quality to the CFA data, whilst the visualization of distal vessels at later PLDs is improved (blue arrow). B) Perfusion images reconstructed from the same raw dataset demonstrate much stronger perfusion signal at later PLDs than the CFA data (green arrow).

The benefit of VFA over CFA is highlighted in Figure 5. Temporal mean images demonstrate the improved distal vessel visibility for angiography and the higher apparent SNR for perfusion imaging. The split-scan repeatability metrics quantitatively demonstrate an improvement in signal stability for VFA compared with CFA, particularly at later PLDs, which was significant (p < 0.001) for both angiography and perfusion imaging: an increase in repeatability of up to 84% was found for angiography (at PLD = 974 ms) and up to 68% for perfusion imaging (at PLD = 1456 ms).

**Figure 5:**
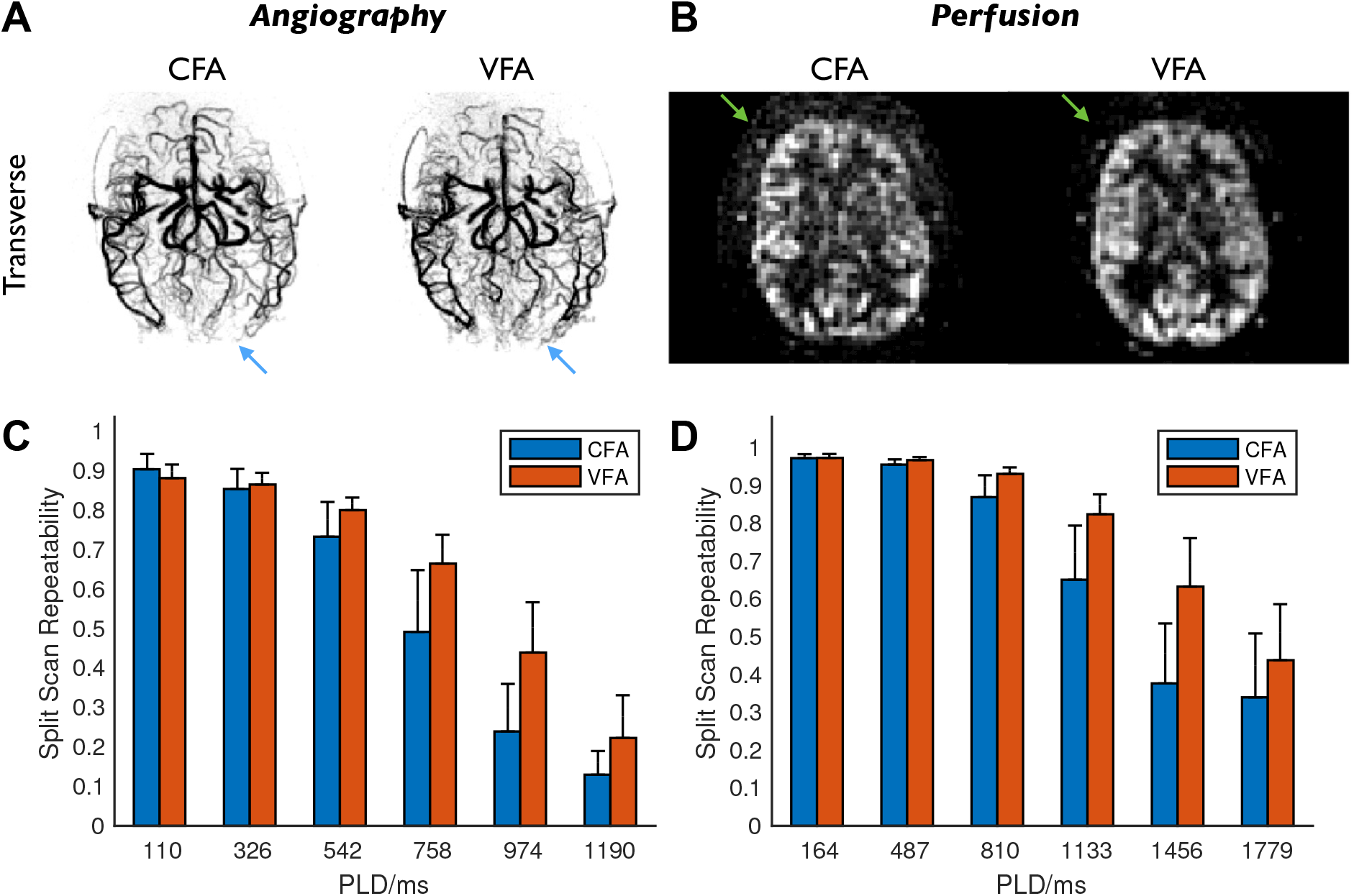
Comparison of CFA and VFA protocols: A) temporal mean angiographic transverse maximum intensity projections of subject 1, showing comparable image quality, with slightly better distal vessel visibility in the VFA protocol (blue arrows); B) temporal mean of transverse perfusion images with PLDs greater than 1 s in subject 1, showing reduced noise in the VFA data (green arrows); C) split scan repeatability (correlation coefficients) for angiography, showing the mean and standard deviation across subjects at each of the first six PLDs; D) split scan repeatability for perfusion imaging at all PLDs. CFA and VFA images in A) and B) are scaled separately for improved visualization. CFA and VFA differences are significant (p < 0.001) for both angiography and perfusion imaging.

### Comparison of reconstruction approaches

A comparison of SENSE and LLR approaches for VFA angiographic reconstructions is shown in Figure 6. Whilst SENSE gives a reasonably clear visualization of the vessels at early timepoints, there is some residual background noise which increases over time, likely due to increased physiological noise from the higher static tissue signal associated with the VFA schedule, obscuring some smaller distal vessels (Figure 6A, inset). In contrast, the LLR approach reduces a lot of this background noise, giving a much clearer depiction of the arterial system, especially the distal vessels at later timepoints.

**Figure 6:**
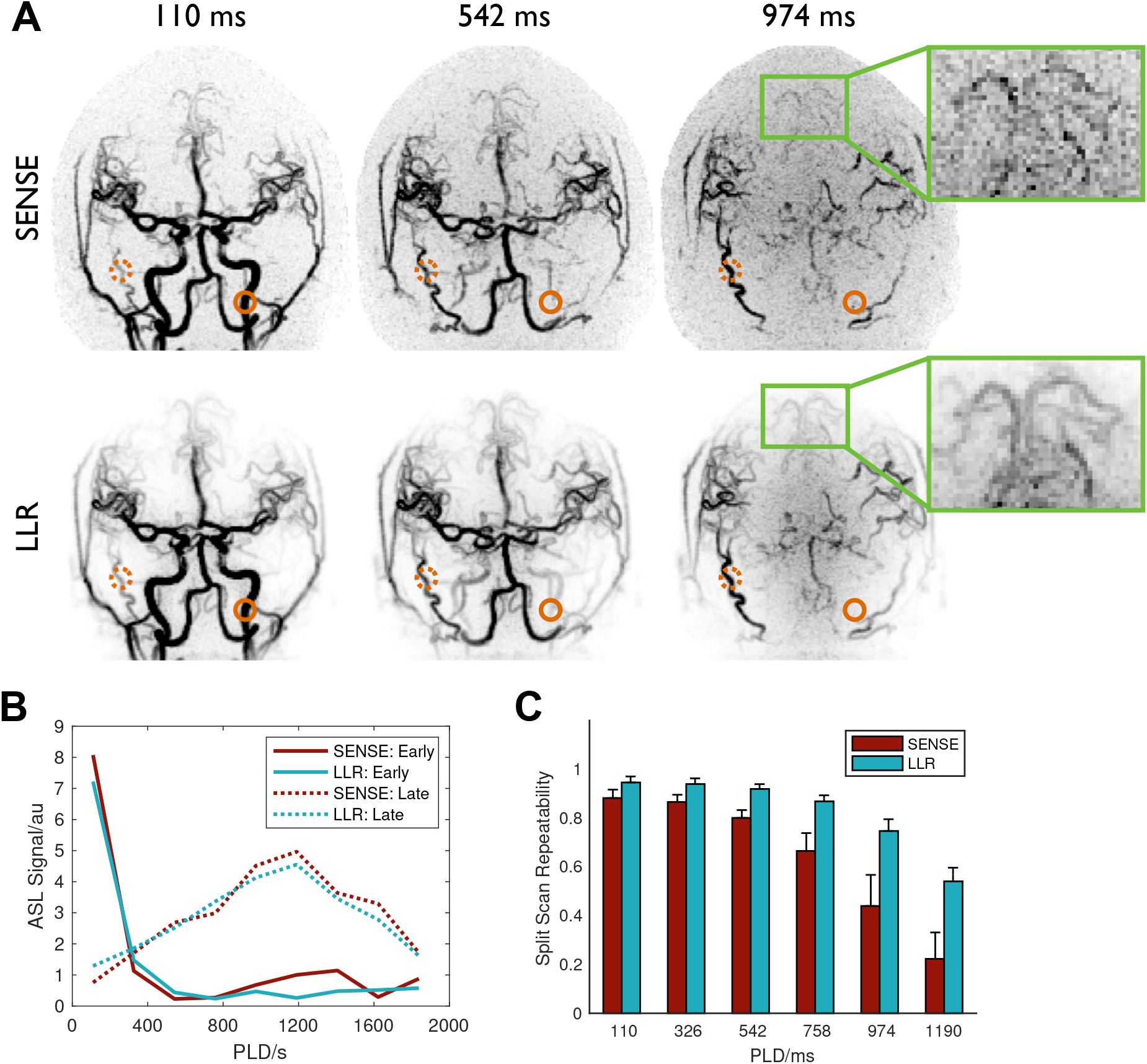
Comparison of angiographic image reconstruction approaches applied to VFA data: A) coronal MIPs of selected frames from SENSE (top row) and LLR (bottom row) reconstructions in subject 2, with the inset showing a zoomed and re-windowed region highlighting the denoising effect of the LLR reconstruction that improves distal vessel visibility; B) signal timeseries from example voxels with early (solid lines) and late (dotted lines) blood arrival (highlighted with circles in A), demonstrating the minimal temporal bias introduced by using the LLR reconstruction approach; C) mean and standard deviation split scan repeatability across all subjects for the first six PLDs showing the significant (p < 0.001) improvement in signal stability achievable using the LLR method, particularly at later time points.

One potential concern with reconstruction approaches such as LLR is that overregularization could bias the signal time courses. Figure 6B shows the angiographic signal in two example voxels with early and late blood arrival, demonstrating that the LLR approach has a denoising effect but does not strongly distort the signal evolution.

LLR also significantly improves the split scan repeatability metric (Figure 6C, p < 0.001), implying improved signal stability, especially at later PLDs. An increase in repeatability of up to 143% relative to SENSE was observed (at PLD = 1190 ms).

Figure 7 compares SENSE and LLR for VFA perfusion imaging. The improvement in image quality achievable with LLR is even more apparent in this case, particularly the denoising effect on the later PLD images when the signal is mostly perfusion weighted (with minimal macrovascular signal). As before, the LLR reconstruction does not appear to strongly bias the signal timecourses (Figure 7B) and gives a significant (p < 0.001) improvement in split scan repeatability (up to 103% at PLD = 1779 ms).

**Figure 7:**
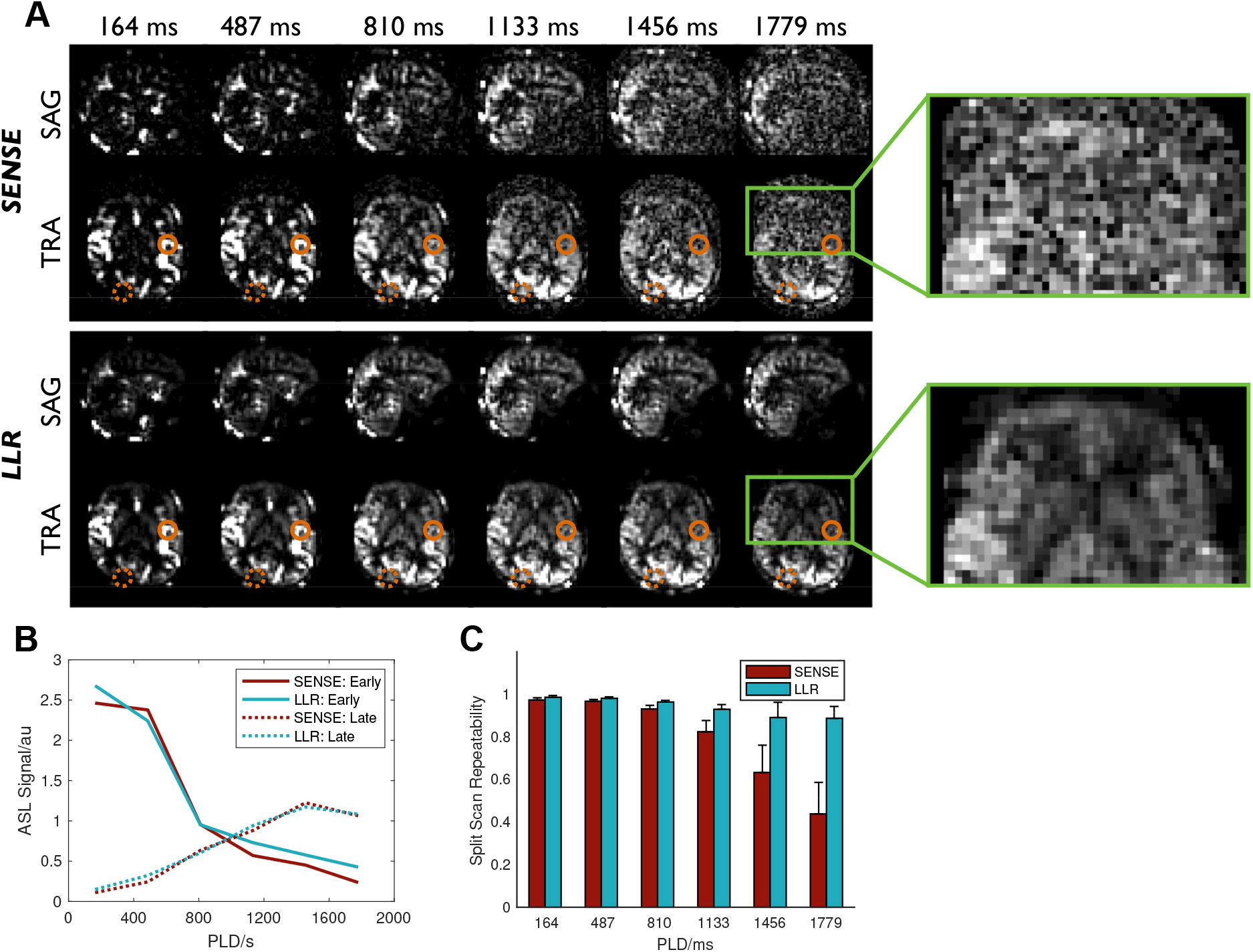
Comparison of perfusion image reconstruction approaches applied to VFA data: A) sagittal (SAG) and transverse (TRA) slices at all PLDs from SENSE (top row) and LLR (bottom row) reconstructions in subject 3, with the inset showing a zoomed and re-windowed region highlighting the noise reduction achieved with the LLR reconstruction; B) signal timeseries from example voxels with early (solid lines) and late (dotted lines) blood arrival (highlighted with circles in A), demonstrating the minimal temporal bias introduced by using the LLR reconstruction approach; C) mean and standard deviation split scan reproducibility across all subjects for all PLDs showing the significant (p < 0.001) improvement in signal stability achievable using the LLR method, particularly at later time points.

LLR angiographic and perfusion reconstructions across multiple timepoints for all subjects can be seen in Supporting Information Videos S1 and S2, respectively, demonstrating the consistent image quality that was obtained in all cases.

## Discussion

In this study we have demonstrated the extension of CAPRIA to a 4D technique, allowing time-resolved 3D angiograms and perfusion images to be obtained from a single non-contrast acquisition. We further optimized the image quality using a variable flip angle readout scheme to reduce the signal attenuation at early timepoints and boost the weaker signal at later timepoints. We also leveraged the spatiotemporal correlations in the data using a locally low rank reconstruction method to significantly reduce the noise level, giving a much clearer delineation of distal vessels and the weaker perfusion signal at long PLDs.

### Pulse sequence design

The extension of the original golden ratio approach^12^ to allow sampling in 3D k-space^22^ was ideal for 4D CAPRIA, enabling flexibility in the temporal and spatial resolution of the reconstruction to allow either angiographic or perfusion-like images to be reconstructed at any PLD within the readout window. Despite the high undersampling factors (R = 19), the sparse and high SNR nature of the angiographic signal allowed reconstruction with minimal apparent artifacts due to the incoherent aliasing that arises from this trajectory, as previously noted for similar acquisition schemes^23,24,42^. The flexibility to retrospectively choose the spatiotemporal resolution allowed perfusion images to be reconstructed with acceleration factors close to one, which was important for this much lower SNR and less sparse signal.

In this study, only a single temporal resolution was demonstrated for angiography and perfusion imaging reconstructions, although any temporal window up to a maximum value, *t*_*max*_, could be chosen, as was demonstrated previously^13^. The *t*_*max*_ approach has the advantage that data can be retrospectively divided into different sections acquired during different time periods (e.g. the first and second half of the scan) which are exactly equivalent to performing shorter scans with higher acceleration factors. This feature could be useful for discarding data acquired during severe motion, for example. However, for temporal windows much smaller or larger than *t*_*max*_ the trajectory will stray further from the near-ideal golden ratio ordering, resulting in reduced SNR efficiency and worse undersampling artifacts. An alternative approach that maintains exact golden ratio ordering for any temporal window^43^ has shown promise for application in combined angiography and perfusion imaging^14^. This method could be easily adapted for use with the 3D radial trajectory used in this study and will be explored in future work.

The use of a relatively long labeling duration (1.4 s) meant that the dynamic angiograms mostly show the outflow of labeled blood water, which might be less intuitive to interpret than the inflow of labeled blood that can be visualized with pulsed ASL techniques^43,44^. However, inflow can be visualized from this kind of data using an inflow subtraction technique^45–47^, as shown in Supporting Information Video S3. However, this reduces SNR and does not easily account for voxels with delayed blood arrival (*δ*_*t*_ > *τ*) or for the potential increase in signal over time resulting from the use of a VFA schedule (see Figure 2B). An alternative approach is to fit a kinetic model and then simulate images at arbitrary timepoints that would be produced with an infinitely long labeling duration, with the effects of RF attenuation and T1 decay removed^26^.

### Flip angle schedules

The quadratic variable flip angle (VFA) scheme used in this study gives an additional degree of freedom in the pulse sequence design, allowing reduced signal attenuation at early timepoints whilst boosting the ASL signal at later timepoints. Simulations and *in vivo* experiments both confirm that this approach is highly beneficial to boost the lower SNR perfusion signal, particularly because of its later arrival in the tissue. Although the angiographic signal is considerably reduced when using small initial flip angles, this did not seem to cause an obvious loss in image quality. This suggests the angiographic signal already has a sufficiently high SNR and the apparent “noise” in these images is dominated by aliased signal and physiological noise (which both scale with signal strength) rather than thermal noise. In contrast, the perfusion signal, which is much weaker due to blood dispersal and the greater degree of T1 decay by the time the labeled water exchanges into tissue, clearly benefits from the increase in signal achieved with the VFA scheme at later timepoints.

An arbitrary weighting factor of ½ was chosen to upweight the perfusion contribution in the flip angle optimization. Other factors could also be chosen, based on the relative importance of angiographic and perfusion information for any given application. Whilst we chose to focus on CFA and quadratic^25^ VFA schedules in this work, other VFA schemes could also be explored, such as linear variations^48^ or the use of a recursive formula to maintain the ASL signal at a constant level^7,49,50^. The use of a balanced steady-state free precession readout could lead to a significant increase in SNR efficiency^44^ and might also benefit from a VFA schedule, although a separate optimization would need to be performed to account for the refocusing of transverse magnetization. However, this approach can suffer from significant signal loss when using a large field of view (FOV, e.g. the whole head) at 3T due to B_0_ inhomogeneity^45^.

### Reconstruction approaches

The locally low-rank (LLR) reconstruction method was found to give a significant improvement in image quality for both angiography and perfusion imaging compared with conventional SENSE. This approach exploits the very strong correlations between the signal timecourses across voxels within any given patch of the image (e.g. they typically have similar arrival times and signal dispersion), meaning that the signal can be well represented by a low rank model. Therefore, enforcing a locally low rank solution that is consistent with the measured data suppresses noise and residual aliasing very effectively. The variation in the k-space trajectory across timepoints combined with a spatiotemporal regularized reconstruction like LLR can also be thought of as reducing the effective undersampling factor of the reconstruction by sharing information across time, thereby reducing noise amplification, which has been shown to benefit similar techniques in ASL perfusion imaging^51^ and angiography^34^. However, it is clear that such regularized approaches have the potential to bias the signal, both spatially and temporally, which could have implications for image interpretation and kinetic model fitting. The regularization factors used here were chosen to minimize this issue, although the optimum choice may be application dependent, and the interaction with model fitting needs to be explored in future work.

Although the image reconstruction comparison in this study focused on VFA data, a similar benefit is likely to be gained in using a LLR reconstruction on CFA data also. In our preliminary experiments we found LLR to provide more robust results than other compressed-sensing reconstruction approaches, so we focused on that technique here. However, other techniques have been applied to ASL angiography and perfusion imaging, such as wavelets/total variation^52^, magnitude subtraction sparsity^53^, spatial sparsity with temporal smoothness^34^, total generalized variation^51^ and low rank + sparse with non-local filters^54^. A comprehensive comparison is beyond the scope of the current work but would be interesting to explore in the future.

### Comparison to related work

Other approaches for combined angiography and perfusion imaging with ASL have also been proposed. One used time-encoded^55^ PCASL in combination with two readout modules, which were separately optimized for angiographic and perfusion information^11^. Another combined time-encoded PCASL with a 2D multi-slice golden ratio readout^14^, which was utilized to vary both the temporal and spatial resolution of the reconstructions. The use of time-encoding allows the generation of images at different effective delay times after the labeling period, with short delays used for angiography and longer delays for perfusion imaging, whilst minimizing the number of excitation pulses required after each ASL preparation. This reduces the attenuation of the ASL signal compared to the proposed approach, where temporal information is obtained only from the use of a long readout, and was shown to significantly increase SNR, especially for the perfusion phase.

However, the use of time-encoding required the effective bolus duration to be short for angiography, which could reduce angiographic SNR. In addition, the relative timings of the images are linked to the PLD associated with each Hadamard encoding block, providing less flexibility than the continuous golden ratio approach used in this study. Finally, the same k-space spokes had to be acquired many times to accommodate all the Hadamard encoding steps, which increased the undersampling factor relative to conventional PCASL, which would be particularly problematic for a 3D radial trajectory like the one proposed in this study.

Compared to multi-slice 2D methods, a 3D golden ratio readout has the additional advantages of SNR efficiency and the ability to reconstruct signals in all voxels across all timepoints. Another option is a 3D stack-of-stars radial trajectory^43,53^, which uses in-plane radial and through-plane Cartesian sampling. This approach may have some advantages when a limited number of slices are required, although the ability to undersample in the through-slice direction is perhaps more limited. However, the optimal combination of time-encoding and readout approach is not yet clear and may be somewhat application dependent, so will be the subject of future investigations.

### Limitations

This study had a number of limitations which could be improved upon in future work. Firstly, the use of non-selective excitation means that the labeling plane is included in the imaging FOV. The different saturation effects on the static tissue at the labeling plane in label and control conditions creates a large difference signal which aliases into the brain, degrading image quality. In addition, a very large reconstructed FOV is necessary to encompass the sensitive area of the receive coil, which increases the computational burden of the reconstruction. A slab-selective excitation could therefore be beneficial. In addition, methods to minimize dephasing effects in proximal vessels, such as flow-compensation or a reduced TE, would be desirable.

Secondly, the magnetization preparation could be improved: only a pre-saturation module was used for background suppression. Additional inversion pulses could reduce physiological noise and improve image quality^56^. The PCASL pulse train duration could also be extended to improve signal strength, but must be balanced against the cost of increased imaging time or undersampling factor. The scan time of 10 minutes used here may already be too long to fit into a busy clinical protocol, so further acceleration is desirable.

Finally, we aimed here to improve qualitative angiography and perfusion imaging, but the extraction of quantitative physiological parameters would be useful. Adaptations to the physiological models described here to include dispersion would be necessary for this, as well as procedures for signal calibration, and will be investigated in the near future.

## Conclusions

4D CAPRIA provides time-resolved angiographic and perfusion information from a single scan across the whole head, with minimal blurring, distortion or dropout artifacts. A quadratic VFA scheme greatly improves image quality at later timepoints, especially for perfusion imaging, and an LLR reconstruction scheme makes further improvements to noise reduction, allowing the clear depiction of distal vessels and late tissue perfusion.

## Acknowledgements

Many thanks to Peter Jezzard and Karla Miller for insightful discussions and support, for the facilities provided by the Oxford Acute Vascular Imaging Centre, and for funding support from the Royal Academy of Engineering (RF/132, RF/201617/16/23) and a Sir Henry Dale Fellowship jointly funded by the Wellcome Trust and the Royal Society (220204/Z/20/Z). The Wellcome Centre for Integrative Neuroimaging is supported by core funding from the Wellcome Trust (203139/Z/16/Z). Many thanks also to Jeff Fessler, Philipp Ehses and colleagues for making available their excellent NUFFT and Siemens raw data reading MATLAB code, as well as to Siemens Healthineers for providing the base pulse sequence code, that we built upon in this work. For the purpose of open access, the author has applied a CC BY public copyright licence to any Author Accepted Manuscript version arising from this submission.

## Data Availability Statement

CAPRIA image reconstruction and signal simulation code used in this study are openly available in GitHub (https://github.com/tomokell/capria_tools) and Zenodo (http://doi.org/10.5281/zenodo.6821643)^57^. Data underlying the plots and code to produce them, including statistical analysis, are openly available in Zenodo (http://doi.org/10.5281/zenodo.6823138)^58^. Deidentified image data will be made available on the WIN Open Data server. This is currently in development. Register here to find out when materials are available for download: https://web.maillist.ox.ac.uk/ox/subscribe/win-open-data.

## Figures

**Supporting Information Figure S1:**
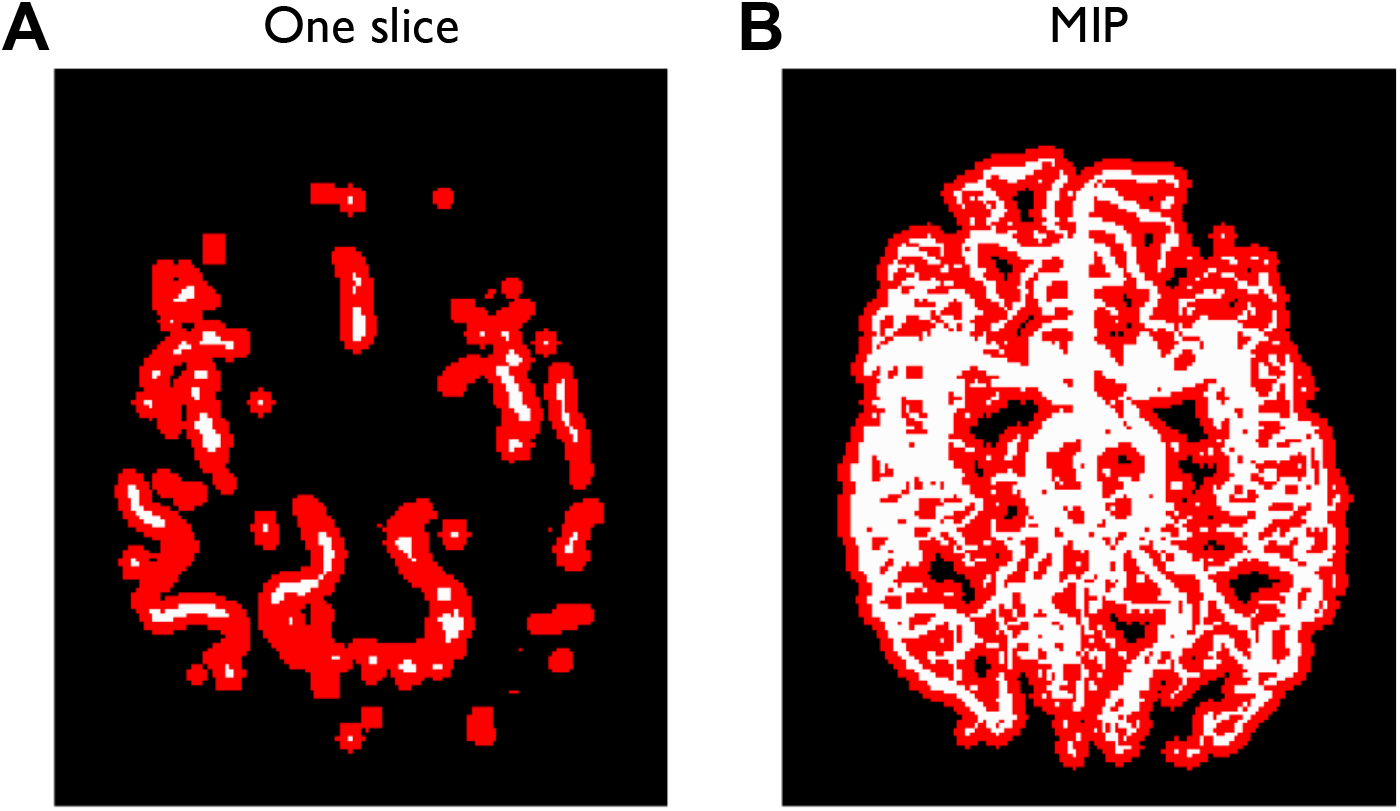
An example vessel mask (white) and dilated vessel mask (red) used for angiography repeatability analysis, shown as a single transverse slice (A) and a transverse maximum intensity projection (B).

Supporting Information Video S1: The first six frames of LLR angiographic reconstructions for all subjects (columns), with MIPs in sagittal (top row), coronal (middle row) and transverse (bottom row) views.

Supporting Information Video S2: All frames of LLR perfusion reconstructions for all subjects (columns), with example sagittal (top row), coronal (middle row) and transverse (bottom row) slices shown.

Supporting Information Video S3: The first six frames of LLR angiographic reconstructions for all subjects (columns), with MIPs in sagittal (top row), coronal (middle row) and transverse (bottom row) views, after inflow subtraction has been applied.

## References

1. Donahue MJ, Achten E, Cogswell PM, et al. Consensus statement on current and emerging methods for the diagnosis and evaluation of cerebrovascular disease. Journal of Cerebral Blood Flow & Metabolism. 2018;38:1391–1417.

2. Suzuki Y, Fujima N, van Osch MJP. Intracranial 3D and 4D MR Angiography Using Arterial Spin Labeling: Technical Considerations. MRMS. 2020;19:294–309.

3. Agarwal R, Brunelli SM, Williams K, Mitchell MD, Feldman HI, Umscheid CA. Gadolinium-based contrast agents and nephrogenic systemic fibrosis: a systematic review and meta-analysis. Nephrol Dial Transplant. 2009;24:856–863.

4. Malayeri AA, Brooks KM, Bryant LH, et al. National Institutes of Health Perspective on Reports of Gadolinium Deposition in the Brain. Journal of the American College of Radiology. 2016;13:237–241.

5. Dixon WT, D. LN, Faul DD, Gado M, Rossnick S. Projection angiograms of blood labeled by adiabatic fast passage. Magn Reson Med. 1986;3:454–462.

6. Nishimura DG, Macovski A, Pauly JM, Conolly SM. MR angiography by selective inversion recovery. Magnetic Resonance in Medicine. 1987;4:193–202.

7. Wang SJ, Nishimura DG, Macovski A. Multiple-readout selective inversion recovery angiography. Magn Reson Med. 1991;17:244–251.

8. Williams DS, Detre JA, Leigh JS, Koretsky AP. Magnetic resonance imaging of perfusion using spin inversion of arterial water. Proceedings of the National Academy of Sciences of the United States of America. 1992;89:212–6.

9. Detre JA, Leigh JS, Williams DS, Koretsky AP. Perfusion imaging. Magn Reson Med. 1992;23:37–45.

10. Alsop DC, Detre JA, Golay X, et al. Recommended implementation of Arterial Spin-Labeled perfusion MRI for clinical applications: A consensus of the ISMRM Perfusion Study group and the European consortium for ASL in dementia. Magnetic Resonance in Medicine. 2015;73:102–116.

11. Suzuki Y, Helle M, Koken P, Van Cauteren M, van Osch MJP. Simultaneous acquisition of perfusion image and dynamic MR angiography using time-encoded pseudo-continuous ASL. Magnetic Resonance in Medicine. 2018;79:2676–2684.

12. Winkelmann S, Schaeffter T, Koehler T, Eggers H, Doessel O. An optimal radial profile order based on the Golden Ratio for time-resolved MRI. IEEE Trans Med Imaging. 2007;26:68–76.

13. Okell TW. Combined angiography and perfusion using radial imaging and arterial spin labeling. Magnetic Resonance in Medicine. 2019;81:182–194.

14. van der Plas Mce, Schmid S, Versluis MJ, Okell TW, Osch MJP van. Time-encoded golden angle radial arterial spin labeling: Simultaneous acquisition of angiography and perfusion data. NMR in Biomedicine. 2021;34:e4519.

15. Okell TW. Combined Angiography and Perfusion using Radial Imaging and Arterial Spin Labeling: Preliminary 4D Results. In: Proceedings 22nd Annual Scientific Meeting, British Chapter of the ISMRM. Leeds, UK; 2016. pp. 34, O5-1.

16. Okell TW. 4D Combined Angiography and Perfusion using Radial Imaging and Arterial Spin Labeling. In: Proceedings 25th Scientific Meeting, ISMRM. Hawaii, USA; 2017. p. 675.

17. Chiew M, Okell TW. Improved Golden Ratio Radial Arterial Spin Labeling Angiography Reconstruction using k-t Sparsity Constraints. In: Proceedings 26th Scientific Meeting, ISMRM. Paris, France; 2018. p. 3351.

18. Golay X, Petersen ET, Hui F. Pulsed star labeling of arterial regions (PULSAR): a robust regional perfusion technique for high field imaging. Magn Reson Med. 2005;53:15–21.

19. Dai W, Garcia D, de Bazelaire C, Alsop DC. Continuous flow-driven inversion for arterial spin labeling using pulsed radio frequency and gradient fields. Magnetic Resonance in Medicine. 2008;60:1488–1497.

20. Okell TW, Chappell MA, Woolrich MW, Günther M, Feinberg DA, Jezzard P. Vessel-encoded dynamic magnetic resonance angiography using arterial spin labeling. Magn Reson Med. 2010;64:698–706.

21. Okell TW, Chappell MA, Kelly ME, Jezzard P. Cerebral blood flow quantification using vessel-encoded arterial spin labeling. J Cereb Blood Flow Metab. 2013;33:1716–1724.

22. Chan RW, Ramsay EA, Cunningham CH, Plewes DB. Temporal stability of adaptive 3D radial MRI using multidimensional golden means. Magn Reson Med. 2009;61:354–363.

23. Wu H, Block WF, Turski PA, Mistretta CA, Johnson KM. Noncontrast-enhanced three-dimensional (3D) intracranial MR angiography using pseudocontinuous arterial spin labeling and accelerated 3D radial acquisition. Magn Reson Med. 2013;69:708–715.

24. Koktzoglou I, Meyer JR, Ankenbrandt WJ, et al. Nonenhanced arterial spin labeled carotid MR angiography using three-dimensional radial balanced steady-state free precession imaging. J Magn Reson Imaging. 2015;41:1150–1156.

25. Schmitt P, Speier P, Bi X, Weale P, Mueller E. Non-contrast-enhanced 4D intracranial MR angiography: Optimizations using a variable flip angle approach. In: Proceedings 18th Scientific Meeting, ISMRM. Stockholm, Sweden; 2010. p. 402.

26. Okell TW, Chappell MA, Schulz UG, Jezzard P. A kinetic model for vessel-encoded dynamic angiography with arterial spin labeling. Magnetic Resonance in Medicine. 2012;68:969–979.

27. Buxton RB, Frank LR, Wong EC, Siewert B, Warach S, Edelman RR. A general kinetic model for quantitative perfusion imaging with arterial spin labeling. Magnetic Resonance in Medicine. 1998;40:383–396.

28. Woods JG, Chappell MA, Okell TW. A general framework for optimizing arterial spin labeling MRI experiments. Magnetic Resonance in Medicine. 2019;81:2474–2488.

29. Mugler JP, Brookeman JR. Three-dimensional magnetization-prepared rapid gradient-echo imaging (3D MP RAGE). Magn. Reson. Med. 1990;15:152–157.

30. Fessler JA, Sutton BP. Nonuniform fast fourier transforms using min-max interpolation. IEEE Transactions on Signal Processing. 2003;51:560–574.

31. Fessler J. Image Reconstruction Toolbox. http://web.eecs.umich.edu/~fessler/irt. Published 2015. Accessed March 1, 2015.

32. Walsh DO, Gmitro AF, Marcellin MW. Adaptive reconstruction of phased array MR imagery. Magnetic resonance in medicine. 2000;43:682–90.

33. Buehrer M, Pruessmann KP, Boesiger P, Kozerke S. Array compression for MRI with large coil arrays. Magn. Reson. Med. 2007;57:1131–1139.

34. Schauman SS, Chiew M, Okell TW. Highly accelerated vessel-selective arterial spin labeling angiography using sparsity and smoothness constraints. Magnetic Resonance in Medicine. 2020;83:892–905.

35. Pruessmann KP, Weiger M, Börnert P, Boesiger P. Advances in sensitivity encoding with arbitrary k-space trajectories. Magn. Reson. Med. 2001;46:638–651.

36. Trzasko J, Manduca A, Borisch E. Local versus global low-rank promotion in dynamic MRI series reconstruction. In: Proceedings 19th Scientific Meeting, ISMRM. Montreal; 2011. p. 4371.

37. Coifman RR, Donoho DL. Translation-Invariant De-Noising. In: Antoniadis A, Oppenheim G, editors. Wavelets and Statistics. Vol. 103. Lecture Notes in Statistics. New York, NY: Springer New York; 1995. pp. 125–150.

38. Taylor AB, Hendrickx JM, Glineur F. Exact Worst-Case Performance of First-Order Methods for Composite Convex Optimization. SIAM J. Optim. 2017;27:1283–1313.

39. Jenkinson M, Beckmann CF, Behrens TEJ, Woolrich MW, Smith SM. FSL. NeuroImage. 2012;62:782–790.

40. Jenkinson M. Improved Optimization for the Robust and Accurate Linear Registration and Motion Correction of Brain Images. NeuroImage. 2002;17:825–841.

41. Fisher RA. Frequency Distribution of the Values of the Correlation Coefficient in Samples from an Indefinitely Large Population. Biometrika. 1915;10:507.

42. Berry ESK, Jezzard P, Okell TW. The advantages of radial trajectories for vessel-selective dynamic angiography with arterial spin labeling. Magnetic Resonance Materials in Physics, Biology and Medicine. 2019;32:643–653.

43. Song HK, Yan L, Smith RX, et al. Noncontrast enhanced four-dimensional dynamic MRA with golden angle radial acquisition and K-space weighted image contrast (KWIC) reconstruction. Magn Reson Med. 2014;72:1541–1551.

44. Yan L, Wang S, Zhuo Y, et al. Unenhanced dynamic MR angiography: high spatial and temporal resolution by using true FISP-based spin tagging with alternating radiofrequency. Radiology. 2010;256:270–279.

45. Okell TW, Schmitt P, Bi X, et al. Optimization of 4D vessel-selective arterial spin labeling angiography using balanced steady-state free precession and vessel-encoding. NMR in Biomedicine. 2016;29:776–786.

46. Kopeinigg D, Bammer R. Time-Resolved Angiography using InfLow Subtraction (TRAILS). Magnetic Resonance in Medicine. 2014;72:669–678.

47. Suzuki Y, Okell TW, Fujima N, van Osch MJP. Acceleration of vessel-selective dynamic MR Angiography by pseudocontinuous arterial spin labeling in combination with Acquisition of ConTRol and labEled images in the Same Shot (ACTRESS). Magnetic Resonance in Medicine. 2019;81:2995–3006.

48. Meixner C, Müller M, Schmitter S, et al. Evaluation of a full dynamic density adapted radial 4D-pcASL angiography sequence exploiting B1+-shimming and ramped SPINS excitation at 7T. In: Proceedings 31st Scientific Meeting, ISMRM. London; 2022. p. 208.

49. Woods JG, Schauman SS, Chiew M, Chappell MA, Okell TW. Time-encoded pseudo-continuous arterial spin labeling: increasing SNR in ASL angiography. arXiv. 2022.

50. van der Plas Mce, Teeuwisse WM, Schmid S, Chappell M, van Osch MJP. High temporal resolution arterial spin labeling MRI with whole-brain coverage by combining time-encoding with Look-Locker and simultaneous multi-slice imaging. Magn Reson Med. 2019;81:3734–3744.

51. Spann SM, Shao X, Wang DJJ, et al. Robust single-shot acquisition of high resolution whole brain ASL images by combining time-dependent 2D CAPIRINHA sampling with spatio-temporal TGV reconstruction. NeuroImage. 2020;206:116337.

52. Taso M, Zhao L, Guidon A, Litwiller DV, Alsop DC. Volumetric abdominal perfusion measurement using a pseudo-randomly sampled 3D fast-spin-echo (FSE) arterial spin labeling (ASL) sequence and compressed sensing reconstruction. Magnetic Resonance in Medicine. 2019;82:680–692.

53. Zhou Z, Han F, Yu S, et al. Accelerated noncontrast-enhanced 4-dimensional intracranial MR angiography using golden-angle stack-of-stars trajectory and compressed sensing with magnitude subtraction: Compressed Sensing in Golden-Angle NCE-dMRA. Magn. Reson. Med. 2018;79:867–878.

54. He G, Lu T, Li H, Lu J, Zhu H. Patch tensor decomposition and non-local means filter-based hybrid ASL image denoising. Journal of Neuroscience Methods. 2022;370:109488.

55. Günther M. Highly efficient accelerated acquisition of perfusion inflow series by cycled arterial spin labeling. In: Proceedings of the 16th Annual Meeting of ISMRM. Berlin, Germany; 2007. p. 380.

56. Robson PM, Dai W, Shankaranarayanan A, Rofsky NM, Alsop DC. Time-resolved Vessel-selective Digital Subtraction MR Angiography of the Cerebral Vasculature with Arterial Spin Labeling. Radiology. 2010;257:507–515.

57. Okell, Thomas W, Chiew, Mark. Combined Angiography and Perfusion using Radial Imaging and Arterial Spin Labeling (CAPRIA) Tools: Initial release. Zenodo; 2022. https://zenodo.org/record/6821643.

58. Okell, Thomas W, Chiew, Mark. 4D Combined Angiography and Perfusion using Radial Imaging and Arterial Spin Labeling (CAPRIA): Data and Code to Reproduce Statistics and Figures. Zenodo; 2022. https://zenodo.org/record/6823138.

